# Age at period cessation and trajectories of cardiovascular risk factors across mid and later life: a prospective cohort study

**DOI:** 10.1101/639518

**Authors:** Linda M O’Keeffe, Diana Kuh, Abigail Fraser, Laura D Howe, Debbie A Lawlor, Rebecca Hardy

## Abstract

**What is already known about this topic:** Age at period cessation is associated with cardiovascular disease. Whether age at period cessation adversely affects change in conventional cardiovascular intermediates from mid to later life is not well understood.

**What this study adds:** Women and clinicians concerned about the impact of type and timing of period cessation on conventional cardiovascular intermediates from midlife should be reassured that the impacts over the long term are small.

**Abstract:** *Objective:* To examine the association between age at period cessation (by type of period cessation) and trajectories of anthropometry, blood pressure, lipids and HBA1c from midlife.

*Design:* Prospective cohort study of women recruited to the Medical Research Council National Survey of Health and Development (NSHD).

*Setting:* Population based prospective cohort study.

*Participants:* Women participating in NSHD with a known date of period cessation and at least one measure of each intermediate cardiovascular risk factor.

*Exposures:* Age at period cessation and type of period cessation (hysterectomy compared with natural menopause).

*Outcomes:* Repeated measures of systolic blood pressure, diastolic blood pressure, body mass index (BMI) and waist circumference (WC) from 36 to 69 years and repeated measures of triglyceride, low density lipoprotein cholesterol (LDL-c), high density lipoprotein cholesterol (HDL-c) and glycated haemoglobin (HBA1c) from 53 to 69 years.

*Results:* We found no evidence that age at period cessation was associated with trajectories of log triglyceride, LDL-c and HDL-c from 53 to 69 years and trajectories of blood pressure from 36 to 69 years, regardless of whether period cessation occurred naturally or due to hysterectomy. While we found some evidence of associations of age at period cessation with log BMI, log WC and log HBA1C, patterns were not consistent and differences were small at age 69 years, with confidence intervals that spanned the null. For instance, the difference in log WC at age 69 per year increase in age at natural menopause was 0.003 (95% confidence interval, −0.0002, 0.01) while the difference per year increase in age at hysterectomy was −0.002 (95% CI, −0.005, 0.001).

*Conclusion:* How and when women experience period cessation is unlikely to adversely affect conventional cardiovascular risk factors across mid and later life. Women and clinicians concerned about the impact of type and timing of period cessation on conventional cardiovascular intermediates from midlife should be reassured that the impacts over the long term are small.

## Introduction

A recent systematic review and meta-analysis of observational studies showed that age at period cessation (natural and surgical combined) before 45 years was associated with greater coronary heart disease (CHD) mortality, cardiovascular disease (CVD) mortality but not stroke mortality ^1^. Since that review, several other large studies have supported these findings ^2–4^, with one study suggesting a weaker association for age at surgical menopause than for age at natural menopause ^2^. Evidence for an inverse association of age at natural menopause and cardiovascular risk is also supported by recent findings from a mendelian randomisation (MR) study showing that genetic variants associated with earlier age at natural menopause are associated with increased cardiovascular risk in women ^5^.

In contrast to the large body of literature demonstrating inverse associations between age at period cessation and CVD risk and mortality, the aforementioned systematic review ^1^ highlighted a lack of studies focusing on age at period cessation and intermediate risk factors such as BMI, blood pressure and diabetes. Available longitudinal studies with repeated measures of cardiovascular risk factors across midlife such as the Study of Women’s Health across the Nation (SWAN) and Coronary Artery Risk Development in Young Adults (CARDIA) ^6–8^ focused on acute and immediate changes close to the menopausal transition rather than examining whether age at period cessation was associated with change over the long term ^6–8^. The few available studies that have examined intermediate risk factors have largely demonstrated a lack of association between age at period cessation and body mass index (BMI), waist hip ratio, diastolic blood pressure (DBP), systolic blood pressure (SBP), glucose, risk of diabetes, hypertension and obesity ^9–13^; however, these studies have mostly been cross-sectional comparing different groups of pre-and post-menopausal women, often with only a single measure of the intermediate risk factor and retrospective reports of age at period cessation in women who were already postmenopausal. Thus, prospective analyses of age at period cessation and change in cardiovascular risk factors across midlife are required to clarify the association between age at period cessation and conventional cardiovascular intermediates from midlife to older age; such analyses go beyond examination of age at period cessation and outcomes at a single time point, providing insights into whether trajectories of cardiovascular risk factors from mid to later life differ by age at period cessation.

We examined the association between age at period cessation, by type of period cessation (hysterectomy compared with natural menopause) and five repeated measures of blood pressure, BMI and waist circumference (WC) and from age 36 to 69 years and three measures of lipids and glycated haemoglobin (HBA1c) from age 53 until age 69 years, in the UK Medical Research Council (MRC) National Survey of Health and Development (NSHD).

## Methods

### Participants

The MRC NSHD is a socially stratified sample of 5,362 (2,547 females and 2,815 males) followed 24 times since birth in England, Scotland and Wales in the first week of March 1946 with an additional nine separate postal questionnaires sent to women in midlife ^14,15^. Ethical approval for the most recent visit when participants were aged 69 years was given by Queen Square Research Ethics Committee (13/LO/1073) and Scotland A Research Ethics Committee (14/SS/1009). Participants provided written informed consent for each visit.

### Data

#### Cardiovascular risk factors

##### Blood pressure and anthropometry from 36 to 69 years

SBP, DBP, height, weight and WC were measured at 36, 43, 53, 60-64 and 69 years. Blood pressure was measured at least twice at 36 and 43 years using a Hawksley Random Zero sphygmomanometer ^16^ and three times at ages 53, 60-64 and 69 years using an Omron HEM-705, while the study member was seated and after a short period of rest ^17,18^. Measurements from the Random Zero sphygmomanometer were adjusted using published conversion equations to achieve compatibility with later measurements ^19^. The second blood pressure reading was used unless it was missing, in which case the first was used. Height (cm), weight (kg) and waist (cm) were measured according to a standardised protocol at all home visits and the research clinic. BMI was calculated by dividing weight in kg by height in metres squared.

##### Lipids and HBA1c from 53 to 69 years

Blood based markers (triglycerides, low density lipoprotein cholesterol (LDL-c), high density lipoprotein cholesterol (HDL-c) and HBA1c were measured at 53, 60-64 and 69 years. A venous blood sample was taken by the nurse according to standardised protocol at ages 53 (non-fasting) ^18^, 60-64 (fasting) ^14^ and 69 years (non-fasting) ^15^. Total cholesterol was measured by enzymatic CHOD–PAP. Triglycerides were measured using a glycerol/kinase POD linked reaction of glycerol liberated enzymatically from triglycerides; LDL-c was calculated using the Friedewald formula ^20^; precipitation for measurement of HDL-c was performed using phosphotungstic Mg^2+^. All these measurements were made with a Bayer DAX-72. Samples were analysed for glycosylated haemoglobin with the Tosoh A1C 2.2 Plus Analyser (Tosoh, Tokyo, Japan) using high performance liquid chromatography ^21,22^.

##### Type and timing of menopause

Information on menstrual irregularity, month and year of last menstrual cycle or any operation to remove the uterus or ovaries and monthly hormone replacement therapy (HRT) use was obtained from annual postal questionnaires between ages 47 and 54 years (inclusive) with an additional one at 57 years and from the face to face interviews with nurses at 43, 53 and 60 – 64 years ^23,24^. Age at which periods ceased naturally (defined as a period of at least 12 months without menstruation) or because of bilateral oophorectomy (with or without hysterectomy), or because of hysterectomy with or without unilateral oophorectomy was calculated. We excluded women whose periods stopped for other reasons, such as chemotherapy (n = 37). We also excluded 189 women starting HRT before menopause who had not ceased HRT for at least a year before giving responses about period regularity and the timing of the last period because it was not possible to assign an accurate date of menopause among these women.

##### Potential confounders

We considered the following as potential confounders of the association between age at period cessation and cardiovascular risk factors based on previous analyses in NSHD ^25,26^ and other cohorts ^27,28^; socioeconomic position, parity, HRT use throughout follow-up, age at menarche, smoking and physical activity at age 36 years. BMI at 36 years was considered as an additional confounder for blood pressure, lipids and HBA1c. The 36-year measures were selected for BMI, smoking and physical activity because these measures represent pre-menopausal measures for most women. Further information can be found in eAppendix 1.

##### Sample sizes for analyses

Of the original birth cohort of females (n=2,547), 1,666 women were eligible for inclusion in analyses of blood pressure and anthropometry and 1,563 women were eligible for inclusion in analyses of lipids and HBA1c. eFigure 1 provides an overview of the study design. Of these women, only participants with a known date of period cessation, at least one measure of the risk factor and complete data on all confounders were included in analyses leading to 908/915 women for inclusion in analyses of blood pressure and anthropometry respectively and 787 women for inclusion of analyses of lipids and HBA1c. eTable 1 shows the number of measures available at each time point.

##### Modelling of age-related change in risk factors

We used multilevel models to examine change over time in cardiovascular risk factors ^29^. Multilevel models estimate mean trajectories of the outcome (here cardiovascular risk factors) while accounting for the non-independence (i.e. clustering) of repeated measurements within individuals, change in scale and variance of measures over time and differences in the number and timing of measurements between individuals (using all available data from all eligible participants under a Missing at Random (MAR) assumption that the value of the missing risk factor can be predicted by other measured variables ^30,31^. Changes in triglyceride, LDL-c, HDL-c and HBA1c were modelled using a linear age term (two levels: measurement occasion and individual), allowing risk factors to change linearly from 53 to 69 years. Linear spline multilevel models (two levels: measurement occasion and individual) were used to model change in SBP, DBP, BMI and WC. Linear splines allow knot points to be fitted in order to derive different periods in which change is approximately linear. In this analysis, we fit a knot point at age 53 years resulting in two periods of change for SBP, DBP, BMI and WC from 36 to 53 years and from 53 to 69 years. All trajectories were modelled in MLwiN version 2.36 ^32^, called from Stata version 14 ^33^ using the runmlwin command.^34^

In all models, age (in years) was centred at the first available measure (ie. at age 36 for SBP, DBP, BMI and WC and at age 53 for lipids and HBA1c). Values of cardiovascular risk factors that had a skewed distribution (BMI, WC, triglyceride, HBA1c) were (natural) log transformed prior to analyses. Further details of model selection are provided in Supplementary Material (eAppendix 2) and model fit statistics are provided in eTables 2 & 3.

##### Association of age at period cessation and type of period cessation with change in risk factors

We examined the association of age at period cessation and type of period cessation (hysterectomy compared to natural menopause) with trajectories of each risk factor. We included variables for type of period cessation and age at period cessation and their interaction with age. We also included an interaction term for age at period cessation and type of period cessation to examine whether the association of age at period cessation differed by type of period cessation. Age at period cessation was centred at the mean age of age at period cessation for the sample (age 50 years).

##### Additional and sensitivity analyses

We examined the characteristics of participants included in analyses of anthropometry compared with those excluded due to missing exposure, outcome or confounder data or loss-to follow-up to better understand the role of selection bias. We examined whether pharmacologic treatment of blood pressure, lipids and HBA1c could have influenced our findings by adding a range of constant values to the risk factor measurements of any individual reporting being on treatment at each time point (see eAppendix 3 for further details).

In order to examine whether findings differed by type of hysterectomy, we tested whether the associations of age at period cessation with cardiovascular risk factors differed between women who had hysterectomy with bilateral oophorectomy compared with hysterectomy with conservation of at least one ovary. Trajectories of risk factors were examined in women who could have been pre- or post-menopausal at age 53. Therefore, we performed a sensitivity analysis excluding women who were still pre-menopausal at 53 years, to understand if our findings differed when the analyses were restricted to post-menopausal measures of cardiovascular risk factors.

## Results

Of the 908/915 women included in analyses of blood pressure and anthropometry, most women were pre-menopausal at age 36 years (99.6% of women who had a natural menopause and 89.6% of women who had a hysterectomy), when the first measure of these was available. Of the 787 women included in analyses of lipids and HBA1c, 94% of women having a hysterectomy had the surgery by 53 years and 61% of women who underwent a natural menopause were post-menopausal by age 53 years. Women who had a hysterectomy were more likely to be in a lower social class, have higher parity, higher prevalence of current smoking at age 36, higher prevalence of physical inactivity at age 36, higher prevalence of HRT use across all time points and lower mean age at menarche and age at period cessation compared with women who had a natural menopause (Table 1).

**Table 1.** Characteristics of included in primary analyses of anthropometry by menopause type (N=915)

|  | **Natural Menopause**<br>n= 675 (n=672 (99.6% pre-menopausal at age 36<br>years (1<sup>st</sup> measure of<br>BMI/WC) | **Hysterectomy**<br>n=240 (n=215 (89.6% pre-menopausal at age 36<br>years 1<sup>st</sup> measure of<br>BMI/WC) |
| --- | --- | --- |
|  | **n (%)** | **n (%)** |
| **Household social class** |  |  |
| Professional/intermediate | 256 (37.9) | 76 (31.7) |
| Skilled (non-manual) | 241 (35.7) | 80 (33.3) |
| Skilled manual and partly skilled | 132 (19.6) | 70 (29.2) |
| Unskilled | 46 (6.8) | 14 (5.8) |
| **Parity** |  |  |
| 0 | 100(14.8) | 15(6.3) |
| 1 or 2 | 379(56.2) | 130(54.2) |
| 3 or more | 196(29.0) | 95(39.6) |
| **Current smoking at age 36** | 233 (34.5) | 91 (37.9) |
| **Physical activity at age 36** |  |  |
| Inactive | 275 (40.7) | 110 (45.8) |
| Less active | 167 (24.7) | 53 (22.1) |
| Most active | 233 (34.5) | 77 (32.1) |
| **HRT use** |  |  |
| Age 36 | 4/652 (0.6) | 5/235 (2.1) |
| Age 43 | 15/623(2.4) | 17/228 (7.5) |
| Age 53 | 91/512 (17.8) | 101/174 (58.1) |
| Age 60-64 | 9/498 (1.8) | 22/ 173 (12.7) |
| Age 69 | 8/465 (1.7) | 8/168 (4.8) |
| **Mean age at menarche (SD)** | 13.1 (1.3) | 12.9 (1.1) |
| **Mean age at period cessation (SD)** | 51.5 (3.9) | 44.1 (6.2) |

BMI, body mass index; HRT, hormone replacement therapy; SD, standard deviation; WC, waist circumference.

Women included in analyses were likely to be in a higher social class, have lower smoking prevalence at age 36, lower HRT use at from 53 to 69 and have undergone a hysterectomy compared to women excluded from analyses due to missing exposure, confounder and outcome data. However, women included in analyses did not differ on parity, physical activity, mean age at menarche, mean age at period cessation, mean BMI at age 36 or mean SBP at age 36 compared to women excluded from analyses (eTable 4).

### Age at period cessation and cardiovascular risk factors

#### Associations with blood pressure and anthropometry from age 36 years

Age at period cessation was also not associated with SBP and DBP at age 36 and change in these from 36 to 69 years in unadjusted (eTable 5) and confounder adjusted analyses (eTable 6 and Figure 1 & 2). In unadjusted analyses (eTable 5), a one-year older age at natural menopause was associated with higher log WC at age 36 years albeit with confidence intervals that spanned the null value; there was no evidence that older age at natural menopause was associated with change in WC from 36 to 69 years and the difference at age 36 persisted at age 69 years, albeit with confidence intervals that spanned the null value. Conversely, a one-year older age at hysterectomy was associated with lower log WC at 36 and lower log WC at age 69 years, though with confidence intervals that spanned the null value. The same patterns of association were observed between age at period cessation and log BMI at ages 53 and 69 years, although associations were weaker. Findings for log WC and log BMI were similar in confounder adjusted analyses (eTable 6 and Figure 1 & 2).

**Figure 1.**
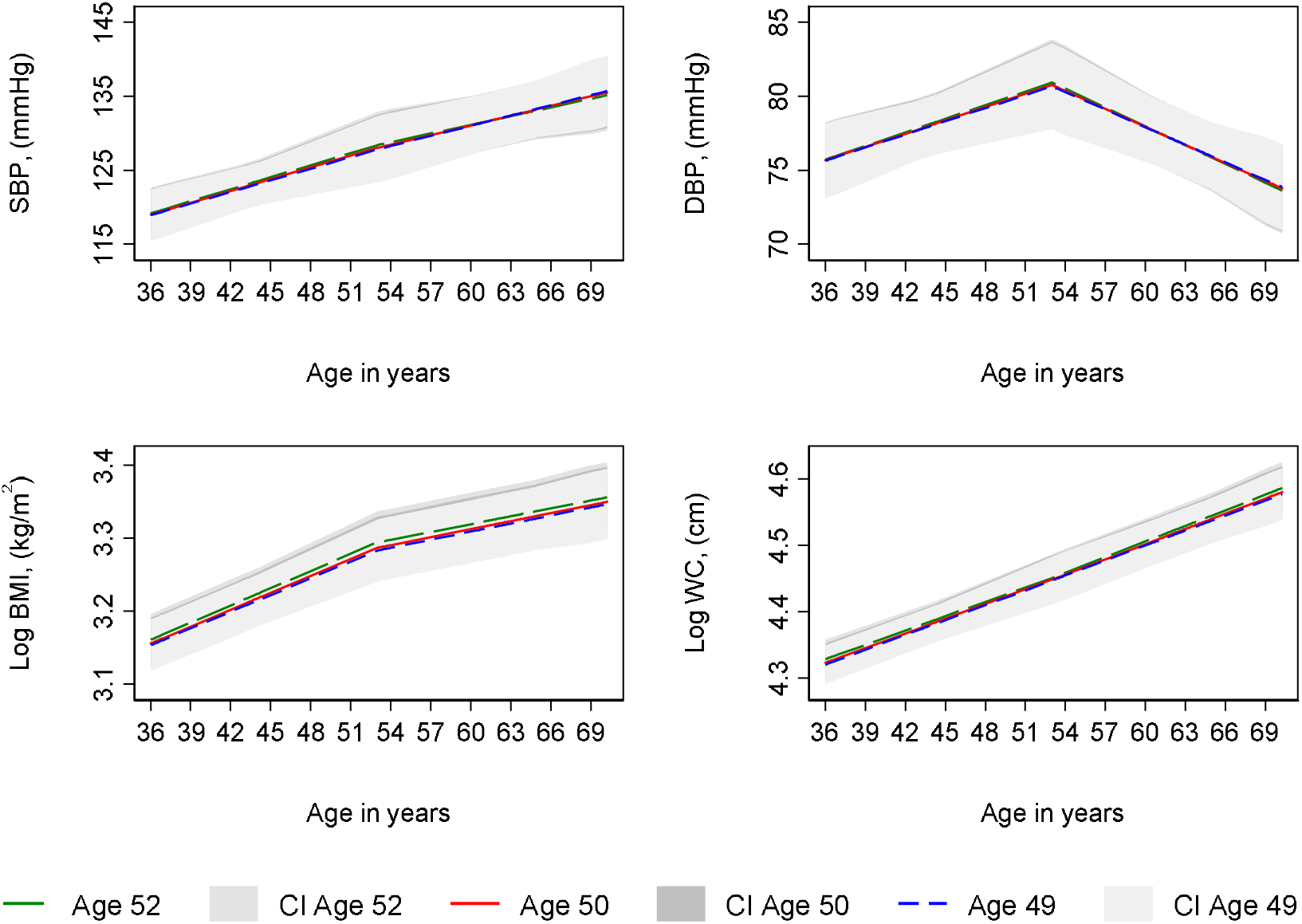
Mean predicted confounder adjusted trajectories of SBP, DBP, log BMI and log waist circumference from 36 to 69 years, by age of at natural menopause. **Legend:** DBP, diastolic blood pressure; SBP, systolic blood pressure; WC, waist circumference. BMI and WC are natural log transformed. Trajectories adjusted for socioeconomic position, type of period cessation, parity, time-varying hormone replacement therapy use, age at menarche, BMI at age 36 (SBP and DBP only), smoking at age 36, physical activity at age 36. Trajectories for the 75^th^ (age 52, green line), median (age50, red line) and 25^th^ percentile(age 49, blue line) of age at period cessation among women with a natural menopause.

**Figure 2.**
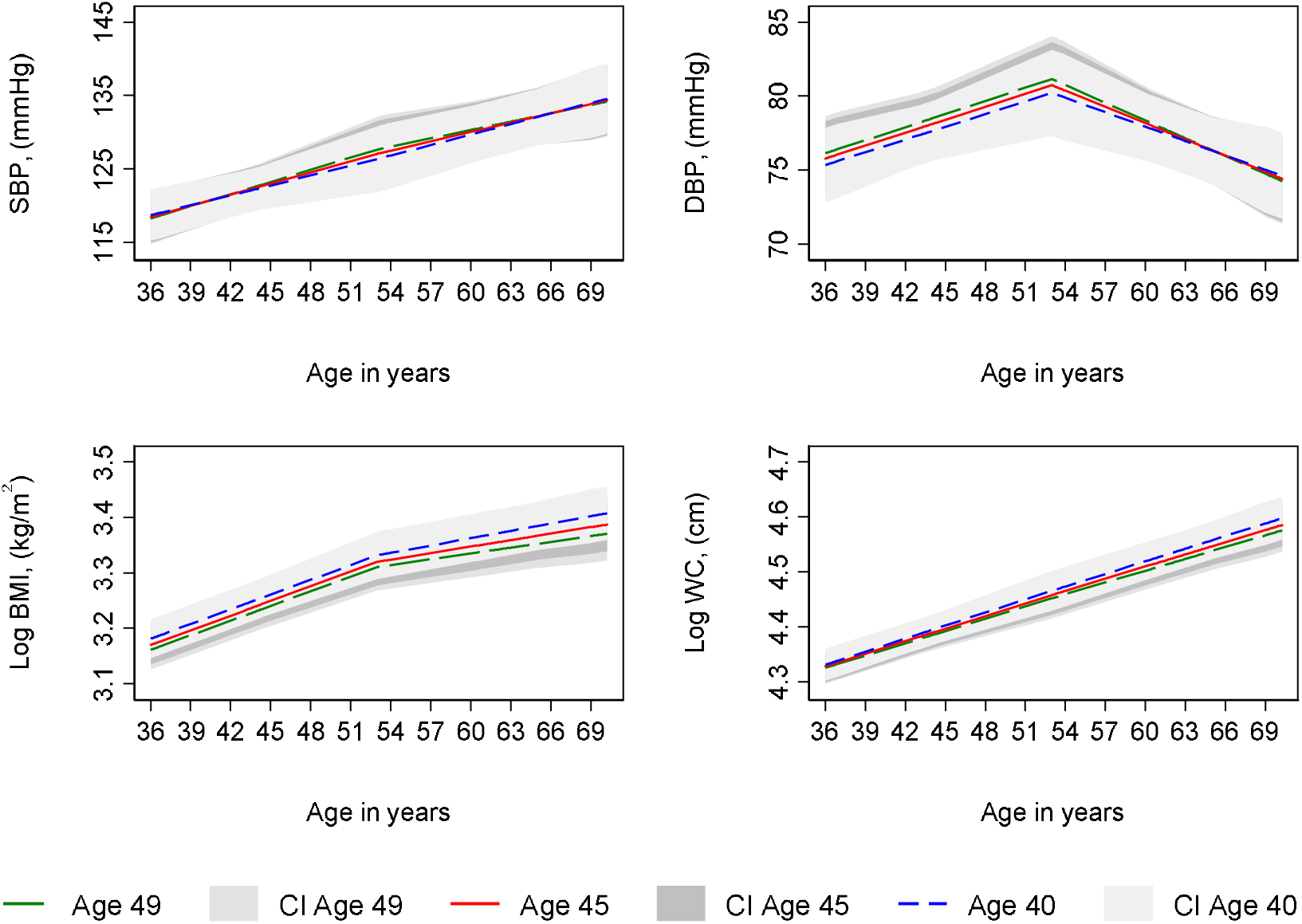
Mean predicted confounder adjusted trajectories of SBP, DBP, log BMI and log waist circumference from 36 to 69 years, by age at hysterectomy. **Legend:** DBP, diastolic blood pressure; SBP, systolic blood pressure; WC, waist circumference. Trajectories adjusted for socioeconomic position, type of period cessation, parity, time-varying hormone replacement therapy use, age at menarche, BMI at age 36 (SBP and DBP only), smoking at age 36, physical activity at age 36. Trajectories for the 75^th^ (age 49, green line), median (age45, red line) and 25^th^ percentile(age 40, blue line) of age at period cessation among women with a natural menopause.

#### Association with lipids and HBA1c from age 53 years

Age at period cessation, whether due to natural menopause or hysterectomy, was not associated with log triglyceride, LDL-c and HDL-c from at age 53 and change in these from 53 to 69 years in unadjusted (eTable 7) or confounder adjusted analyses (eTable 8 and Figure 3 & 4). In unadjusted analyses (eTable 7), age at natural menopause and age at hysterectomy were each associated with lower log HBA1c at age 53 years, albeit with confidence intervals that spanned the null value. However, a one-year older age at natural menopause was associated with a faster increase in log HBA1c from 53 to 69 years such that by 69 years, older menopause was associated with higher log HbA1c, although confidence intervals spanned the null value. In contrast, older age at hysterectomy was associated with slightly slower increases in log HBA1c from 53 to 69 leading to lower log HBA1c at age 69 years, albeit with a confidence interval that spanned the null value. Findings for log HBA1c were similar in confounder adjusted analyses (eTable 8 and Figure 3 & 4).

**Figure 3.**
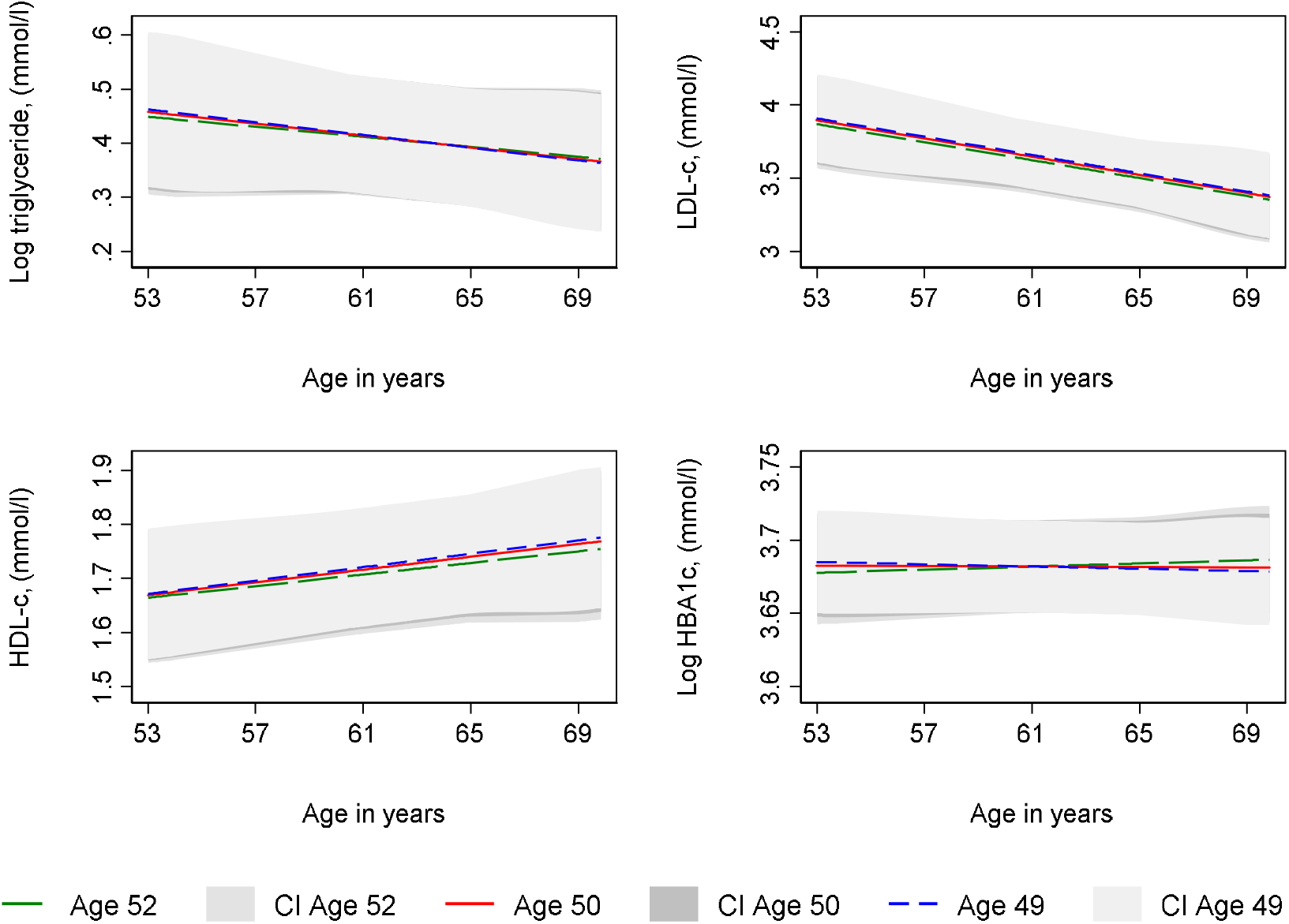
Mean predicted confounder adjusted trajectories of lipids and HBA1c from 53 to 69 years, by age at natural menopause. **Legend:** HBA1c, glycated haemoglobin; HDL-c, high density lipoprotein cholesterol; LDL-c, low density lipoprotein cholesterol. HBA1c and triglyceride are natural log transformed. Trajectories adjusted for socioeconomic position, type of period cessation, parity, time-varying hormone replacement therapy use, age at menarche, BMI at age 36 (SBP and DBP only), smoking at age 36, physical activity at age 36. Trajectories for the 75^th^ (age 52, green line), median (age50, red line) and 25^th^ percentile(age 49, blue line) of age at period cessation among women with a natural menopause.

**Figure 4.**
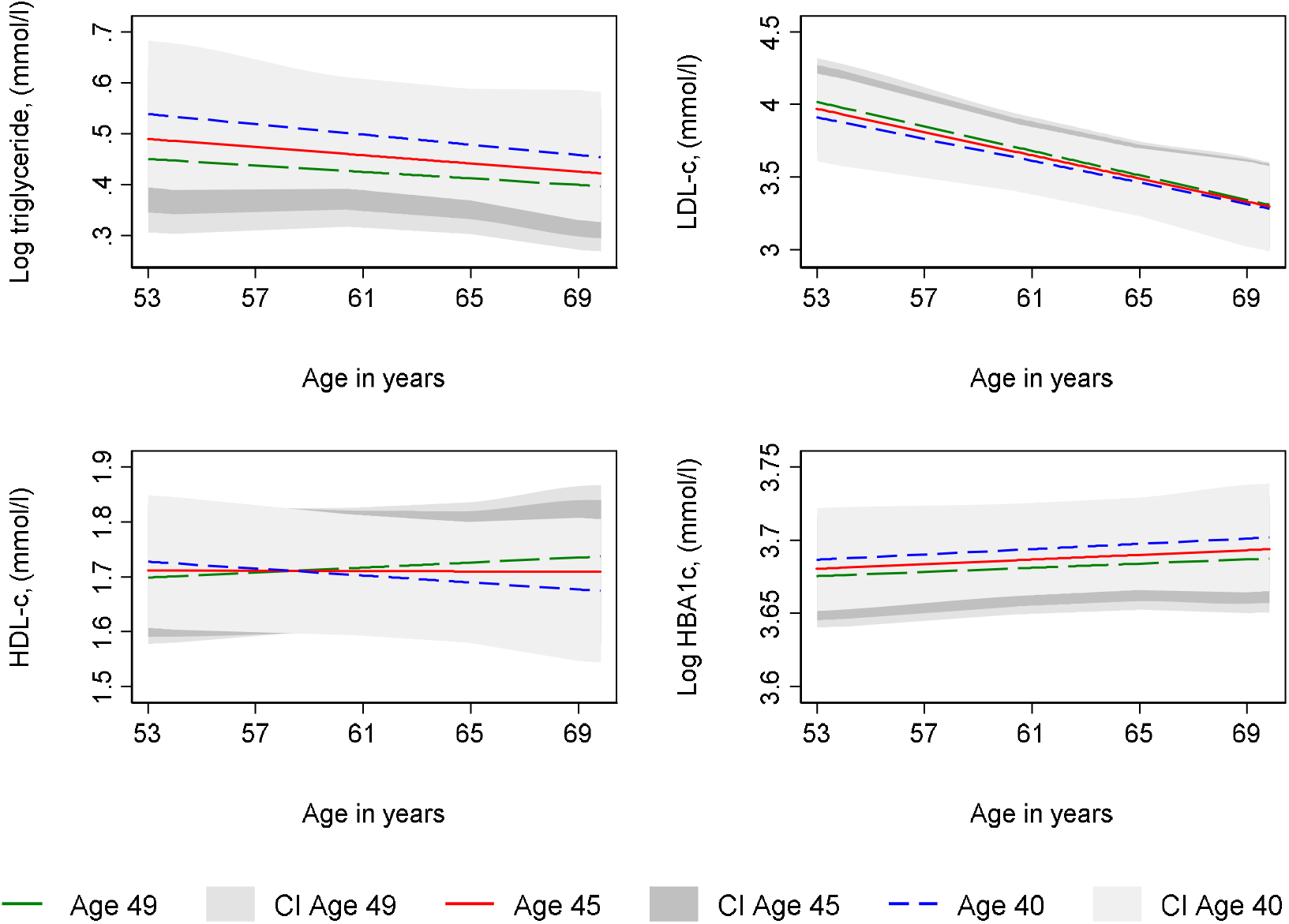
Mean predicted confounder adjusted trajectories of lipids and HBA1c from 53 to 69 years, by age at hysterectomy. **Legend:** HBA1c, glycated haemoglobin; HDL-c, high density lipoprotein cholesterol; LDL-c, low density lipoprotein cholesterol. HBA1c and triglyceride are natural log transformed. Trajectories adjusted for socioeconomic position, type of period cessation, parity, time-varying hormone replacement therapy use, age at menarche, BMI at age 36 (SBP and DBP only), smoking at age 36, physical activity at age 36. Trajectories for the 75^th^ (age 49, green line), median (age45, red line) and 25^th^ percentile(age 40, blue line) of age at period cessation among women with a natural menopause.

#### Type of period cessation

Type of period cessation (hysterectomy compared with natural menopause) was not strongly associated with trajectories of SBP, DBP, log BMI and log WC from 36 to 69 years (eTable 9 and Figure 5) in unadjusted and confounder adjusted analyses or with log triglyceride and LDL-c from 53 to 69 years (eTable 10 and Figure 6). In unadjusted analyses, there was some evidence of faster increases in log HBA1c and faster decreases in HDL-c from 53 to 69 years among women who had a hysterectomy (eTable 10); however, these differences attenuated upon adjustment for confounders (eTable 10 and Figure 6).

**Figure 5.**
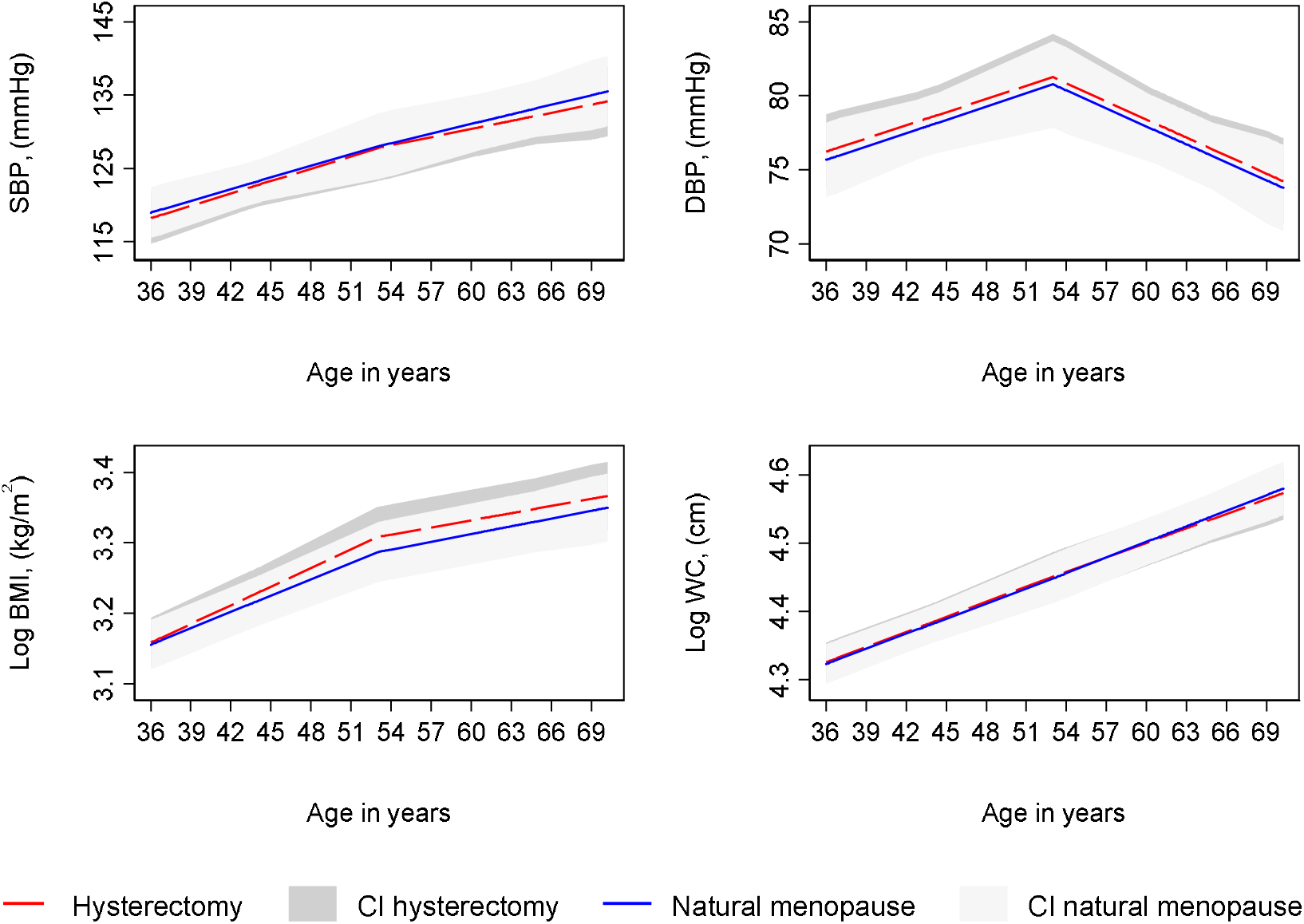
Mean predicted confounder adjusted trajectories of SBP, DBP, log BMI and log waist circumference from 36 to 69 years, by type of period cessation. **Legend:** DBP, diastolic blood pressure; SBP, systolic blood pressure; WC, waist circumference. Trajectories adjusted for socioeconomic position, age at period cessation, parity, time-varying hormone replacement therapy use, age at menarche, BMI at age 36 (SBP and DBP only), smoking at age 36, physical activity at age 36.

**Figure 6.**
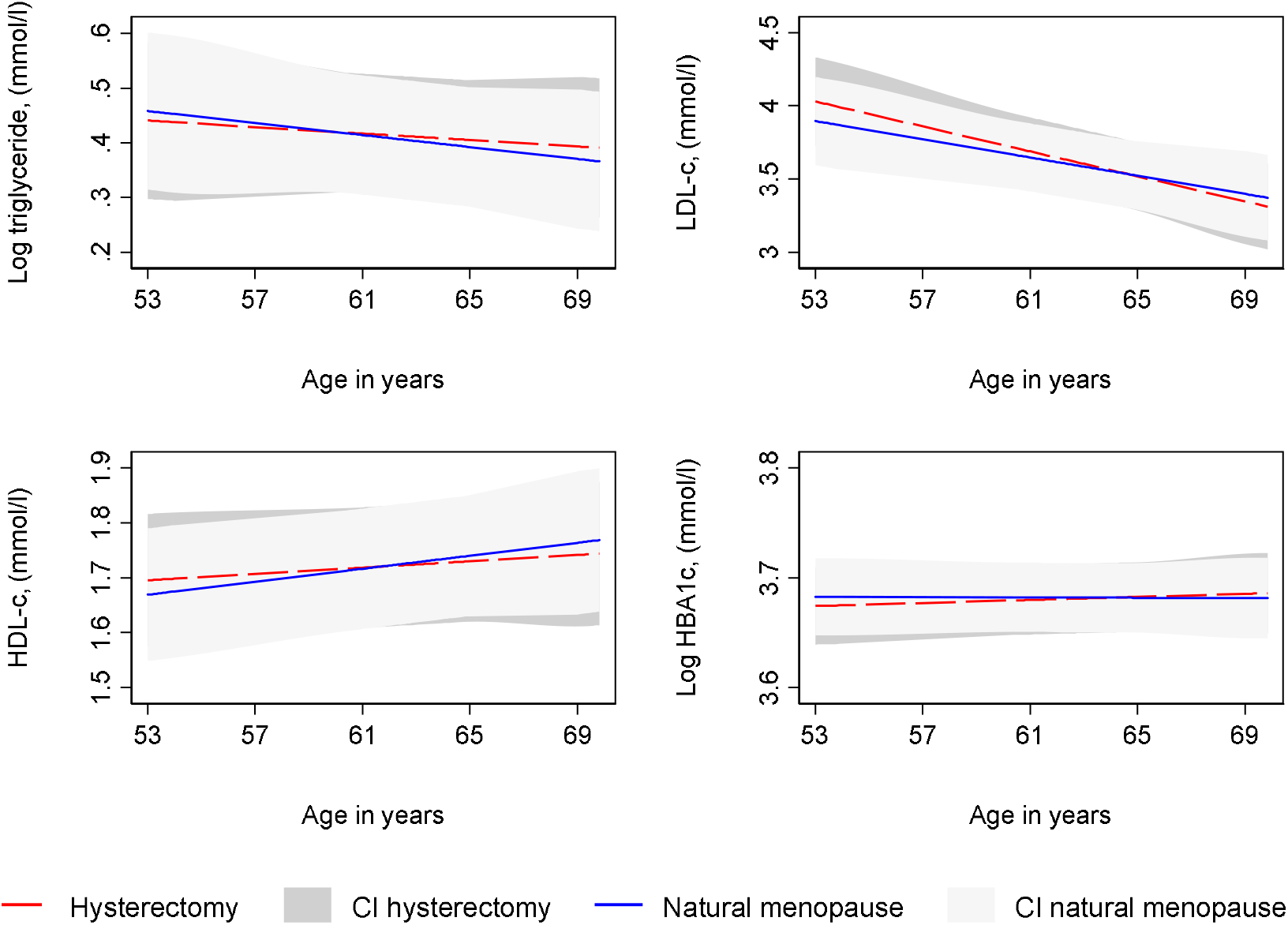
Mean predicted confounder adjusted trajectories of lipids and glycated haemoglobin from 53 to 69 years, by type of period cessation. **Legend:** HBA1c, glycated haemoglobin; HDL-c, high density lipoprotein cholesterol; LDL-c, low density lipoprotein cholesterol. HBA1c and triglyceride are natural log transformed. Trajectories adjusted for socioeconomic position, age at period cessation, parity, time-varying hormone replacement therapy use, age at menarche, BMI at age 36 (SBP and DBP only), smoking at age 36, physical activity at age 36.

#### Additional and sensitivity analyses

We found little evidence that the association of age at hysterectomy with any of the cardiovascular risk factors varied by type of procedure (conservation of at least one ovary or bilateral oophorectomy). Our findings were not altered when we accounted for treatment of lipids, HBA1c and blood pressure. In sensitivity analyses excluding women who had reached menopause after 53 years, findings for the association of age at period cessation or type of period cessation with cardiovascular risk factors were not appreciably altered compared with our primary analyses; all these results are available on request from the authors.

## Discussion

In a British birth cohort with prospective assessment of age at period cessation and repeated measures of eight conventional cardiovascular intermediates from 36/53 to 69 years, we found little evidence of associations between age at period cessation or type of period cessation and trajectories of anthropometry and blood pressure (from 36 years) and lipids and HBA1c (from 53 years) up to age 69 years.

### Comparison with other studies

A lack of studies focusing on intermediate cardiovascular risk factors was highlighted in the most recent systematic review of age at period cessation and CVD ^1^. Our findings are comparable with the few available studies of intermediate cardiovascular risk factors included in that review. For instance, in a cross-sectional study of Chinese women aged 40 to 59 years (N=2,498 postmenopausal women and N=2245 pre-menopausal women), age at natural menopause was not associated with any intermediate risk factors studied including risk of hypertension, diabetes, dyslipidaemia and obesity or mean differences in SBP, DBP, BMI, total cholesterol, triglyceride, HDL-c, LDL-c and glucose ^12^. In the Japan Nurses’ Health Study, another cross-sectional study of 22,426 women, age at natural menopause (categorised as <45, 45 to 53 and >53) was not associated with risk of hypertension or diabetes but early menopause was associated with risk of hypercholesterolemia ^35^, which contrasts with our findings that showed no strong evidence of associations of age at period cessation with lipids in later life. Our findings showing that type of period cessation (hysterectomy compared with natural menopause) was not associated with cardiovascular risk factors are also comparable to findings from the SWAN ^7^ and CARDIA ^8^ studies, which were of similar design to ours and included repeated measures of risk factors across the menopausal transition. However, our study examines change across a wider age range and into the 7^th^ decade of life, demonstrating that type of period cessation is not associated with cardiovascular risk factors once all women have passed through menopause and reach older ages where cardiovascular ill-health is more common.

### Implications

If associations of age at period cessation and CVD events are causal and not the result of confounding in observational studies ^1^ or shared genetic architecture of age at period cessation and CVD ^5^, our findings would suggest that changes in conventional cardiovascular intermediates over the long term are an unlikely mediating pathway. The findings also have important implications for women and clinicians, as they suggest that any impact of age at period cessation or type of period cessation on conventional cardiovascular intermediates over the long term is likely to be small. However, there may still be associations with other cardiovascular intermediates that have not been evaluated here such as coronary artery calcification and vascular structure and function, given that both available studies included in the aforementioned systematic review and meta-analysis showed that older age at period cessation was associated with reduced risk of carotid atherosclerosis ^1^.

Our findings should be considered in the context of the acute but transient changes in lipids observed close to the time of menopause in previous cohorts such as SWAN ^6^ and the Women’s Midlife Health Project ^36^. Although neither of these cohorts directly examined age at period cessation and change in cardiovascular risk factors, it is possible that age at period cessation is associated with incident CVD risk not through a long-term effect on cardiovascular risk factors but through acute effects on risk, which eventually attenuate over time. These acute effects close to the time of menopause may be clinically relevant for later CVD risk. Further work examining age at period cessation and other cardiovascular intermediates such as vascular structure and function may help to further elucidate the mechanisms underlying age at period cessation and CVD risk. Longitudinal cohorts examining age at period cessation and acute effects on intermediate risk factors close to the menopausal transition will also provide a greater understanding of the aetiology of age at period cessation and CVD events, if reported associations in observational studies ^1^ are causal.

### Strengths and Limitations

There are several strengths to our study including the prospective, detailed and longitudinal collection of data on menopausal characteristics and cardiovascular measures from 36 and 53 years. Our study is the first to our knowledge to examine the association of type and timing of period cessation with cardiovascular risk factors into the 7^th^ decade of life. We have included women who have undergone hysterectomy and who were taking HRT, which many previous analyses have excluded, and we have captured the full range of ages at which menopause occurs, which is a limitation of previous analyses that have often excluded women with very early or very late ages at period cessation. We have used multilevel models which take account of clustering of repeated measures within individuals and the correlation between measures over time and performed several sensitivity analyses to examine the robustness of our findings to the effect of pharmacological treatment of risk factors. Limitations include combining non-fasting and fasting bloods for risk factors and the availability of measures from 36 years for only four out of the eight risk factors as data for lipids and HBA1c were only available from age 53 years. Selection bias is also a potential limitation and individuals included in our analysis were more advantaged than those excluded due to missing exposure, outcome and confounder data thus limiting the potential generalisability of our findings to the wider population.

### Conclusion

How and when women experience period cessation is unlikely to adversely affect conventional cardiovascular intermediates from midlife. Women or clinicians concerned about the impact of type and timing of period cessation on conventional cardiovascular intermediates from midlife should be reassured that the impacts over the long term are likely to be small.

## Contributor and guarantor statement

LMOK had the idea for the study, performed all analyses and wrote the manuscript up for publication. All other authors advised on the analysis and provided critical revisions to the manuscript.

## Public and patient involvement

Public and patient involvement was not part of this research.

## Data sharing

Data are available upon submission and approval of a research proposal. Further information can be found at https://www.nshd.mrc.ac.uk/data/.

## Acknowledgements

We thank NSHD study members for their lifelong participation and past and present members of the NSHD study team who helped to collect the data

## References

1. Muka T, Oliver-Williams C, Kunutsor S, et al. Association of age at onset of menopause and time since onset of menopause with cardiovascular outcomes, intermediate vascular traits, and all-cause mortality: a systematic review and meta-analysis. JAMA Cardiology. 2016;1(7):767–776.

2. Peters SA, Woodward M. Women’s reproductive factors and incident cardiovascular disease in the UK Biobank. Heart (British Cardiac Society). 2018;104(13):1069–1075.

3. Yang L, Lin L, Kartsonaki C, et al. Menopause Characteristics, Total Reproductive Years, and Risk of Cardiovascular Disease Among Chinese Women. Circulation: Cardiovascular Quality and Outcomes. 2017;10(11):e004235.

4. Ley SH, Li Y, Tobias DK, et al. Duration of reproductive life span, age at menarche, and age at menopause are associated with risk of cardiovascular disease in women. Journal of the American Heart Association. 2017;6(11):e006713.

5. Sarnowski C, Kavousi M, Isaacs S, et al. Genetic variants associated with earlier age at menopause increase the risk of cardiovascular events in women. Menopause. 2018;25(4):451–457.

6. Matthews KA, Crawford SL, Chae CU, et al. Are changes in cardiovascular disease risk factors in midlife women due to chronological aging or to the menopausal transition? Journal of the American College of Cardiology. 2009;54(25):2366–2373.

7. Matthews KA, Gibson CJ, El Khoudary SR, Thurston RC. Changes in cardiovascular risk factors by hysterectomy status with and without oophorectomy: Study of Women’s Health Across the Nation. Journal of the American College of Cardiology. 2013;62(3):191–200.

8. Appiah D, Schreiner PJ, Bower JK, Sternfeld B, Lewis CE, Wellons MF. Is Surgical Menopause Associated With Future Levels of Cardiovascular Risk Factor Independent of Antecedent Levels? The CARDIA Study. American journal of epidemiology. 2015:kwv162.

9. Cho GJ, Lee JH, Park HT, et al. Postmenopausal status according to years since menopause as an independent risk factor for the metabolic syndrome. Menopause. 2008;15(3):524–529.

10. Brand JS, Van Der Schouw YT, Onland-Moret NC, et al. Age at menopause, reproductive life span, and type 2 diabetes risk: results from the EPIC-InterAct study. Diabetes care. 2013;36(4): 1012–1019.

11. Qiu C, Chen H, Wen J, et al. Associations between age at menarche and menopause with cardiovascular disease, diabetes, and osteoporosis in Chinese women. The Journal of Clinical Endocrinology & Metabolism. 2013;98(4):1612–1621.

12. He L, Tang X, Li N, et al. Menopause with cardiovascular disease and its risk factors among rural Chinese women in Beijing: a population-based study. Maturitas. 2012;72(2):132–138.

13. Wu X, Cai H, Kallianpur A, et al. Age at menarche and natural menopause and number of reproductive years in association with mortality: results from a median follow-up of 11.2 years among 31,955 naturally menopausal Chinese women. PloS one. 2014;9(8):e103673.

14. Kuh D, Pierce M, Adams J, et al. Cohort profile: updating the cohort profile for the MRC National Survey of Health and Development: a new clinic-based data collection for ageing research. International journal of epidemiology. 2011;40(1):e1–e9.

15. Kuh D, Wong A, Shah I, et al. The MRC National Survey of Health and Development reaches age 70: maintaining participation at older ages in a birth cohort study. European journal of epidemiology. 2016;31(11):1135–1147.

16. Wills AK, Lawlor DA, Matthews FE, et al. Life course trajectories of systolic blood pressure using longitudinal data from eight UK cohorts. PLoS medicine. 2011;8(6):e1000440.

17. Strand BH, Murray ET, Guralnik J, Hardy R, Kuh D. Childhood social class and adult adiposity and blood-pressure trajectories 36–53 years: gender-specific results from a British birth cohort. Journal of epidemiology and community health. 2010:jech. 2010.115220.

18. Masi S, D’Aiuto F, Martin-Ruiz C, et al. Rate of telomere shortening and cardiovascular damage: a longitudinal study in the 1946 British Birth Cohort. European heart journal. 2014:ehu226.

19. Stang A, Moebus S, Möhlenkamp S, et al. Algorithms for converting random-zero to automated oscillometric blood pressure values, and vice versa. American journal of epidemiology. 2006;164(1):85–94.

20. Skidmore PM, Hardy RJ, Kuh DJ, Langenberg C, Wadsworth ME. Birth weight and lipids in a national birth cohort study. Arteriosclerosis, thrombosis, and vascular biology. 2004;24(3):588–594.

21. Prynne C, Mander A, Wadsworth M, Stephen A. Diet and glycosylated haemoglobin in the 1946 British birth cohort. European Journal of Clinical Nutrition. 2009;63(9):1084–1090.

22. Silverwood RJ, Richards M, Pierce M, et al. Cognitive and kidney function: results from a British birth cohort reaching retirement age. PloS one. 2014;9(1):e86743.

23. Kuh D, Langenberg C, Hardy R, et al. Cardiovascular risk at age 53 years in relation to the menopause transition and use of hormone replacement therapy: a prospective British birth cohort study. BJOG: An International Journal of Obstetrics & Gynaecology. 2005;112(4):476–485.

24. Mishra G, Kok H, Ecob R, Cooper R, Hardy R, Kuh D. Cessation of hormone replacement therapy after reports of adverse findings from randomized controlled trials: evidence from a British birth cohort. American Journal of Public Health. 2006;96(7):1219–1225.

25. Hardy R, Kuh D, Wadsworth M. Smoking, body mass index, socioeconomic status and the menopausal transition in a British national cohort. International journal of epidemiology. 2000;29(5):845–851.

26. Hardy R, Kuh D. Reproductive characteristics and the age at inception of the perimenopause in a British National Cohort. American journal of epidemiology. 1999;149(7):612–620.

27. Willett W, Stampfer MJ, Bain C, et al. Cigarette smoking, relative weight, and menopause. American journal of epidemiology. 1983;117(6):651–658.

28. Bromberger JT, Matthews KA, Kuller LH, Wing RR, Meilahn EN, Plantinga P. Prospective study of the determinants of age at menopause. American journal of epidemiology. 1997; 145(2): 124–133.

29. Goldstein H. Multilevel statistical models; 2nd edition ed. London: Edward Arnold; 1995.

30. Howe LD, Tilling K, Matijasevich A, et al. Linear spline multilevel models for summarising childhood growth trajectories: A guide to their application using examples from five birth cohorts. Statistical Methods in Medical Research. 2013:0962280213503925.

31. Tilling K, Macdonald-Wallis C, Lawlor DA, Hughes RA, Howe LD. Modelling childhood growth using fractional polynomials and linear splines. Annals of Nutrition and Metabolism. 2014;65(2-3): 129–138.

32. MLwiN Version 2.36. Centre for Multilevel Modelling UoBc, 2016 p.

33. Stata 14.0 [computer program]. Texas StataCorp; 2016..

34. runmlwin: Stata module for fitting multilevel models in the MLwiN software. Centre for Multilevel Modelling, University of Bristol [computer program]. 2016.

35. Lee JS, Hayashi K, Mishra G, et al. Independent association between age at natural menopause and hypercholesterolemia, hypertension, and diabetes mellitus: Japan nurses’ health study. Journal of Atherosclerosis. 2012:14746.

36. Do K, Green A, Guthrie J, Dudley E, Burger H, Dennerstein L. Longitudinal study of risk factors for coronary heart disease across the menopausal transition. American journal of epidemiology. 2000;151(6):584–593.

