## Supplementary material for "Age at period cessation and trajectories of cardiovascular risk factors across mid and later life: a prospective cohort study": Online supplemental files

**eAppendix 1 Details on confounders**

We categorised participant’s occupation as reported at age 53 into six classes (from professional to unskilled manual) according to the Registrar General social classification. Parity was self-reported by the participants at all data collections across adulthood and included as a continuous covariate. Women reported monthly histories of HRT use throughout follow-up and from this, HRT use (yes/no) as a time-varying covariate was included in our models. Age at menarche in years was obtained at medical examination by school doctors when participants were 14-15 years old, supplemented for those who had not reached menarche by 15 years (n=94) by retrospective reports of age at menarche by postal questionnaires when participants were aged 48 years ^1^. Smoking, obtained from self-reported information at 36 years, was classified as former/current/never. Physical activity at 36 years, obtained from self-reports of frequency and duration of participation in leisure time activities was classified as inactive (reported no participation); moderately active (participated in relevant activities one to four times: in the previous month); or most active (participated in relevant activities five or more times: in the previous month ^2^.

**eAppendix 2 Details of model selection**

Models were derived by initially examining observed data for each risk factor and plotting mean values for each risk factor over time to examine the possible shape of the trajectory.

Linear spline multilevel models were used for SBP, DBP, WC and BMI. Based on the observed data for these, we compared observed and predicted measurements for a selection of suitable models for each risk factor. We examined rates of change between time periods in order to examine whether changes between periods were similar or different. In cases where rates of change between spline periods appeared identical, the fit of models with reduced splines was explored. Final models for SBP, DBP, BMI and WC had one knot placed at 53 years resulting in two periods of change; from 36-53 and from 53-69. This knot was selected based on examination of observed data over time and comparing model fit statistics (Akaike’s Information Criterion) for several models with different knot points (with knot points placed at whole years closest to mean age at clinics due to a greater density of measures). The selection of this knot point at age 53 years, also had the additional advantage of allowing comparability with the linear slopes modelled from 53 to 69 years for the blood-based biomarkers.

*SBP and DBP*

The models for SBP and DBP took the form of: SBP_ij_ /DBP_ij_ = β_0_ + u_0j_ + (β_1_+ u_1j_ )s_ij1_ + (β_2_+ u_2j_ )s_ij2_ + e_ij_(age_ij_) where for person j at measurement occasion i; β_0_ represents the fixed effect coefficient for the average intercept, β_1_ to β_2_ represent fixed effect coefficients for the average linear slopes of each linear spline, u_0j_ to u_3j_ indicate person-specific random effects for the intercept and slopes respectively, and e_ij_ represents the occasion-specific residuals or measurement error which was allowed to vary with age.

*BMI, WC*

BMI and WC were natural log transformed. The models for BMI and WC took the form of: log BMI_ij_ /log WC_ij_ = β_0_ + u_0j_ + (β_1_+ u_1j_ )s_ij1_ + (β_2_+ u_2j_ )s_ij2_ + e_ij_(age_ij_) where for person j at measurement occasion i; β_0_ represents the fixed effect coefficient for the average intercept, β_1_ to β_3_ represent fixed effect coefficients for the average linear slopes of each linear spline, u_0j_ to u_2j_ indicate person-specific random effects for the intercept and slopes respectively, and e_ij_ represents the occasion-specific residuals or measurement error which was allowed to vary with age.

*Lipids and HBA1c*

Triglyceride and glycated haemoglobin were natural log transformed. Lipids and HBA1c were modelled using a linear age term.The models for lipids and HBA1C took the form of: log triglycerides _ij_ / LDL-c_ij_ /HDL-c_ij /_ log HBA1c _ij_= β_0_ + u_0j_ + (β_1_+ u_1j_) age_ij1_ + e_ij_(age_ij_) where for person j at measurement occasion i; β_0_ represents the fixed effect coefficient for the average intercept, β_1_ represents fixed effect coefficients for the average linear slope, u_0j_ and u_1j_ indicate person-specific random effects for the intercept and slopes respectively, and e_ij_ represents the occasion-specific residuals or measurement error which was allowed to vary with age.

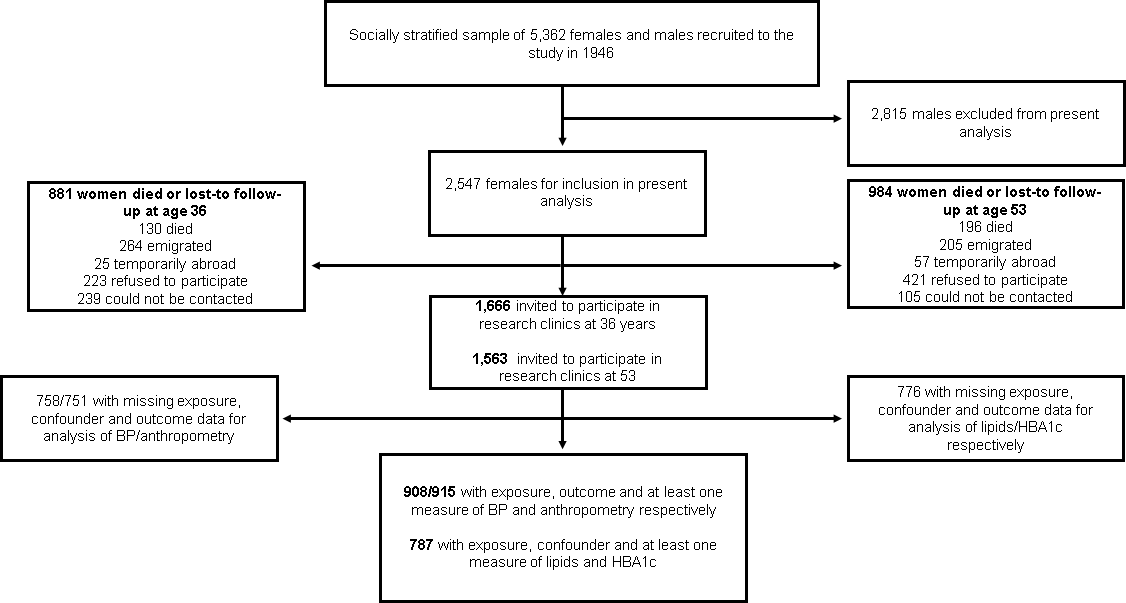

**eFigure 1 Flow diagram of participants included in study**

BP, blood pressure; HBA1c, glycated haemoglobin.

**eTable 1 Number of participants with cardiometabolic measures at each time point**

|  | Age 36 | Age 43 | Age 53 | Age 60-64 | Age 69 | Total measures | | Total participants | | | | Median measures (IQR) |
| --- | --- | --- | --- | --- | --- | --- | --- | --- | --- | --- | --- | --- |
| Lipids/HBA1c† |  |  | 575 | 555 | 520 | 1650 | | | 787 | | 3 (2-3) | |
| BMI/WC | 887 | 858 | 692 | 677 | 637 | 3751 | | | 915 | | 5 (4-5) | |
| SBP/DBP | 887 | 851 | 686 | 671 | 633 | 3728 | | | 908 | | 5 (4-5) | |

DBP, diastolic blood pressure; HBA1C; glycated haemoglobin; HDL-c, high density lipoprotein cholesterol; IQR, interquartile range; LDL-c, low density lipoprotein cholesterol; SBP, systolic blood pressure.

† Triglyceride, HDL-c and LDL-c.

eTable 2 Model details for log triglyceride and log HBA1c trajectories

|  | No of contributing individuals | | Assessment of model fit | | | |
| --- | --- | --- | --- | --- | --- | --- |
|  | Total number of observations | Number of individuals with 1 measure | Mean observed, (SD)* | Mean predicted, (SD)* | Mean difference (observed – predicted) * | 95% level of agreement between observed and predicted* |
| Log triglyceride |  |  |  |  |  |  |
| Overall | 1650 | 787 |  |  |  |  |
| 53 years | 575 | 575 | 0.39 (0.47) | 0.32 (0.29) | 0.0624 | -0.49,0.62 |
| 53 -64 years | 963 | 684 | 0.26 (0.50) | 0.30 (0.28) | -0.0344 | -0.64,0.57 |
| 64-69 years | 686 | 576 | 0.29 (0.46) | 0.24 (0.24) | 0.0474 | -0.58,0.67 |
| Log HBA1c |  |  |  |  |  |  |
| Overall | 1650 | 787 |  |  |  |  |
| 53 years | 575 | 575 | 3.62 (0.14) | 3.63 (0.10) | -0.0054 | -0.12,0.11 |
| 53 -64 years | 963 | 684 | 3.64 (0.13) | 3.64 (0.10) | 0.0032 | -0.10,0.11 |
| 64-69 years | 686 | 576 | 3.67 (0.12) | 3.67 (0.10) | -0.0045 | -0.10,0.09 |
| LDL-c |  |  |  |  |  |  |
| Overall | 1650 | 787 |  |  |  |  |
| 53 years | 575 | 575 | 3.56 (1.01) | 3.66 (0.47) | -0.1000 | -1.46,1.26 |
| 53 -64 years | 963 | 684 | 3.63 (1.03) | 3.58 (0.50) | 0.0582 | -1.32,1.44 |
| 64-69 years | 686 | 576 | 3.19 (1.02) | 3.28 (0.51) | -0.0820 | -1.36,1.19 |
| HDL-c |  |  |  |  |  |  |
| Overall | 1650 | 787 |  |  |  |  |
| 53 years | 575 | 575 | 1.86 (0.48) | 1.84 (0.35) | 0.0159 | -0.35,0.38 |
| 53 -64 years | 963 | 684 | 1.81 (0.46) | 1.82 (0.35) | -0.0062 | -0.36,0.35 |
| 64-69 years | 686 | 576 | 1.77 (0.46) | 1.76 (0.37) | 0.0087 | -0.31,0.33 |

HBA1c, glycated haemoglobin; HDL-c, high density lipoprotein cholesterol; LDL-c, low density lipoprotein cholesterol.

* Triglyceride and HBA1c are natural log transformed. All values are in log form.

**eTable 3 Model details for blood pressure and anthropometry trajectories**

|  | No of contributing individuals | | Assessment of model fit | | | |
| --- | --- | --- | --- | --- | --- | --- |
|  | Total number of observations | Number of individuals with 1 measure | Mean observed, (SD)* | Mean predicted, (SD)* | Mean difference (observed – predicted) * | 95% level of agreement between observed and predicted* |
| SBP |  |  |  |  |  |  |
| Overall | 3728 | 908 |  |  |  |  |
| 36 years | 887 | 887 | 116.90 (13.61) | 115.97 (5.32) | 0.6125 | -19.19,20.41 |
| 36 -53 years | 1739 | 905 | 119.05 (14.95) | 119.29 (7.06) | -0.4514 | -22.10,21.20 |
| 53-69 years | 1989 | 837 | 132.64 (17.85) | 132.37 (9.88) | 0.2714 | -22.44,22.98 |
| DBP |  |  |  |  |  |  |
| Overall | 3728 | 908 |  |  |  |  |
| 36 years | 887 | 887 | 74.77 (11.53) | 74.43 (3.98) | 0.2263 | -17.80,18.25 |
| 36 -53 years | 1739 | 905 | 75.81 (11.81) | 75.90 (4.37) | -0.1485 | -18.57,18.27 |
| 53-69 years | 1989 | 837 | 77.19 (10.92) | 77.08 (5.53) | 0.0956 | -15.68,15.87 |
| Log BMI |  |  |  |  |  |  |
| Overall | 3751 | 915 |  |  |  |  |
| 36 years | 887 | 887 | 3.15 (0.15) | 3.14 (0.14) | 0.0019 | -0.06,0.07 |
| 36 -53 years | 1746 | 912 | 3.18 (0.16) | 3.18 (0.15) | -0.0009 | -0.08,0.08 |
| 53-69 years | 2005 | 844 | 3.32 (0.19) | 3.32 (0.17) | 0.0004 | -0.07,0.07 |
| Log WC |  |  |  |  |  |  |
| Overall | 3751 | 915 |  |  |  |  |
| 36 years | 887 | 887 | 4.33 (0.14) | 4.31 (0.09) | 0.0131 | -0.11,0.14 |
| 36 -53 years | 1746 | 912 | 4.34 (0.14) | 4.34 (0.10) | -0.0060 | -0.14,0.12 |
| 53-69 years | 2003 | 844 | 4.49 (0.15) | 4.49 (0.13) | 0.0043 | -0.10,0.11 |

DBP, diastolic blood pressure; SBP, systolic blood pressure; WC, waist circumference.

*BMI and WC are natural log transformed. All values are in log form.

**eAppendix 3 Details of analyses examining pharmacologic treatment effects**

In order to understand the potential effect of treatment of risk factors on our findings, we performed sensitivity analyses adding 20%, 40% and 60% to the triglyceride values of individuals reporting treatment with lipid lowering medication at each time point, 10%, 20% and 30% to LDL-c values of participants reporting treatment with lipid lowering medication at each time point, and subtracting 5%, 10% and 15% from the HDL-c values of individuals under treatment at each time point. For HBA1c, we performed separate analyses adding 1%, 2% and 3% to the values of individuals reporting treatment with diabetes medications at each time point. Similarly, we examined the potential effect of treatment with antihypertensive medications on our findings, adding 10%, 20% and 30% to the recorded SBP of participants under treatment at each time point, and 5%, 10% and 15% to the recorded DBP of participants under treatment.

eTable 4 Characteristics of women included in primary analyses of anthropometry compared to women excluded from analyses due to missing data

|  | **Included in analysis (N=915)** | **Excluded from analysis (N=448 – 1141)** | **P value** |
| --- | --- | --- | --- |
|  | **n (%)** | **n (%)** |  |
| **Household social class** |  |  |  |
| Skilled (non-manual) | 332 (36.3) | 435 (38.1) | <0.05 |
| Professional/intermediate | 321 (35.1) | 353 (30.9) |  |
| Skilled manual and partly skilled | 198 (22.1) | 302 (26.5) |  |
| Unskilled | 60 (6.6) | 51 (4.5) |  |
| **Parity** |  |  |  |
| 0 | 115 (12.6) | 72 (11.9) | 0.68 |
| 1 or 2 | 509 (55.6) | 347 (57.5) |  |
| 3 or more | 291 (31.8) | 185 (30.6) |  |
| **Current smoking at age 36** | 276 (30.1) | 282 (37.6) | <0.01 |
| **Physical activity at age 36** |  |  |  |
| Inactive | 385 (42.1) | 320 (42.8) | 0.91 |
| Less active | 220 (24.0) | 182 (24.3) |  |
| Most active | 310 (33.9) | 246 (32.9) |  |
| **HRT use** |  |  |  |
| Age 36 | 9 (1.0) | 7 (0.9) | 0.8 |
| Age 43 | 32 (3.7) | 40 (5.2) | 0.15 |
| Age 53 | 193 (27.9) | 239 (46.0) | <0.001 |
| Age 60-64 | 32 (4.7) | 48 (10.4) | <0.001 |
| Age 69 | 16 (2.5) | 22 (4.5) | 0.06 |
| **Type of menopause** |  |  |  |
| Natural | 675 (73.7) | 577 (80.8) | 0.001 |
| Hysterectomy | 240 (26.2) | 137 (19.2) |  |
|  | **Mean (SD)** | **Mean (SD)** |  |
| **Mean age at menarche (SD)** | 13.0 (1.2) | 13.1 (1.1) | 0.35 |
| **Mean age at period cessation (SD)** | 49.6 (5.6) | 49.5 (6.1) | 0.75 |
| **Mean BMI at age 36 (SD)** | 23.6 (3.9) | 23.5 (4.2) | 0.49 |
| **Mean SBP at age 36 (SD)** | 116.9 (13.6) | 17.3 (14.6) | 0.54 |

BMI, body mass index; SBP, systolic blood pressure; SD, standard deviation. Denominator for participants excluded may also vary due to missing data on the characteristics included in the table. *P* value is for the difference in proportions for categorical variables from *χ*² test or difference in means for continuous variables from t tests between included and excluded participants

|  | **Mean trajectory for NM** | **Mean difference in trajectory per year increase in age at NM** | **Mean trajectory for HY** | **Mean difference in trajectory per year increase in HY** | **P value for interaction of type and timing of menopause** |
| --- | --- | --- | --- | --- | --- |
| **Log BMI** |  |  |  |  |  |
| Age 36 | 3.14(3.13,3.15) | 0.002(-0.001,0.005) | 3.15(3.12,3.17) | -0.003(-0.01,0.00001) | 0.13 |
| Δ 36- 53 | 0.01(0.01,0.01) | 0.0001(-0.0001,0.0002) | 0.01(0.01,0.01) | -0.00001(-0.0002,0.0002) | 0.54 |
| Δ 53 -69 | 0.002(0.002,0.003) | -0.00004(-0.0002,0.0001) | 0.002(0.001,0.004) | -0.0001(-0.0003,0.0001) | 0.39 |
| Age 69 | 3.33(3.31,3.34) | 0.002(-0.002,0.007) | 3.35(3.32,3.39) | -0.005(-0.01,-0.0009) | 0.04 |
| **Log WC** |  |  |  |  |  |
| Age 36 | 4.31(4.30,4.32) | 0.002(-0.0003,0.005) | 4.32(4.30,4.34) | -0.001(-0.003,0.002) | 0.20 |
| Δ 36- 53 | 0.007(0.007,0.008) | -0.0001(-0.0002,0.0001) | 0.007(0.006,0.009) | -0.00007(-0.0002,0.0001) | 0.43 |
| Δ 53 -69 | 0.006(0.005,0.007) | 0.0001(-0.0001,0.0003) | 0.006(0.004,0.007) | -0.0001(-0.0002,0.0001) | 0.43 |
| Age 69 | 4.53(4.52,4.55) | 0.002(-0.0010,0.01) | 4.53(4.51,4.56) | -0.003(-0.006,0.0002) | 0.05 |
| **SBP** |  |  |  |  |  |
| Age 36 | 116.0 (114.8,117.2) | 0.02(-0.26,0.30) | 115.1 (112.8,117.5) | -0.06(-0.36,0.24) | 0.91 |
| Δ 36- 53 | 0.90(0.79,1.01) | 0.01(-0.01,0.04) | 1.00(0.78,1.22) | 0.01(-0.01,0.04) | 0.45 |
| Δ 53 -69 | 0.18(0.06,0.30) | -0.03(-0.05,-0.003) | 0.09(-0.14,0.31) | -0.01(-0.04,0.02) | 0.08 |
| Age 69 | 134.1 (132.5,135.8) | -0.22(-0.59,0.14) | 133.5 (130.4,136.6) | -0.02(-0.39,0.35) | 0.30 |
| **DBP** |  |  |  |  |  |
| Age 36 | 74.3(73.4,75.2) | 0.001(-0.20,0.20) | 74.9(73.2,76.6) | 0.06(-0.14,0.27) | 0.84 |
| Δ 36- 53 | 0.41(0.33,0.48) | 0.01(-0.01,0.02) | 0.40(0.25,0.55) | 0.0002(-0.02,0.02) | 0.83 |
| Δ 53 -69 | -0.46(-0.54,-0.38) | -0.01(-0.03,0.01) | -0.46(-0.61,-0.30) | -0.01(-0.03,0.01) | 0.37 |
| Age 69 | 73.9(72.9,74.8) | -0.08(-0.30,0.14) | 74.4(72.5,76.2) | -0.03(-0.26,0.20) | 0.59 |

**eTable 5 Unadjusted association of age at period cessation (per year increase) with anthropometry and blood pressure from 36 to 69 years**

**Legend:** DBP, diastolic blood pressure; log WC, natural log waist circumference; HY, hysterectomy; NM, natural menopause; SBP, systolic blood pressure. Note that BMI and WC are natural log transformed and all values presented are in log form. Δ = change per year in risk factor.

^ P value for the interaction of the association of age at period cessation with type of menopause with trajectories.

eTable 6 Association of age at period cessation (per year increase) with anthropometry and blood pressure from 36 to 69 years, adjusted for co-variates

|  | **Mean trajectory for NM (reference for period cessation at age 50)** | **Mean difference in trajectory per year increase in age at NM** | **Mean trajectory for HY (reference for period cessation at age 50)** | **Mean difference in trajectory per year increase in HY** | **P value for interaction of type and timing of menopause** |
| --- | --- | --- | --- | --- | --- |
| **Log BMI** |  |  |  |  |  |
| Age 36 | 3.16(3.12,3.19) | 0.002(-0.001,0.005) | 3.16(3.11,3.20) | -0.003(-0.005,0.0006) | 0.09 |
| Δ 36- 53 | 0.008(0.006,0.01) | 0.0001(-0.0001,0.0002) | 0.009 (0.006,0.01) | 0.00001(-0.0001,0.0002) | 0.64 |
| Δ 53 -69 | 0.004(0.002,0.006) | -0.00003(-0.0002,0.0001) | 0.003(0.001,0.006) | -0.0001(-0.0003,0.0001) | 0.44 |
| Age 69 | 3.35(3.30,3.40) | 0.003(-0.001,0.007) | 3.36(3.30,3.42) | -0.004(-0.008,-0.0003) | 0.02 |
| **Log WC** |  |  |  |  |  |
| Age 36 | 4.32(4.29,4.35) | 0.003(0.0004,0.005) | 4.33(4.29,4.36) | -0.001(-0.003,0.002) | 0.07 |
| Δ 36- 53 | 0.007(0.005,0.009) | -0.0001(-0.0003,0.0001) | 0.007(0.005,0.010) | -0.00004(-0.0002,0.0001) | 0.46 |
| Δ 53 -69 | 0.008(0.006,0.010) | 0.0001(-0.0001,0.0003) | 0.007(0.004,0.010) | -0.0001(-0.0002,0.0001) | 0.34 |
| Age 69 | 4.57(4.53,4.61) | 0.003(-0.0002,0.01) | 4.56(4.52,4.61) | -0.002(-0.005,0.001) | 0.02 |
| **SBP** |  |  |  |  |  |
| Age 36 | 119.0(115.5,122.4) | 0.08(-0.20,0.36) | 118.2(113.9,122.5) | -0.07(-0.38,0.24) | 0.80 |
| Δ 36- 53 | 0.53(0.21,0.85) | 0.01(-0.02,0.03) | 0.56(0.16,0.96) | 0.01(-0.02,0.04) | 0.63 |
| Δ 53 -69 | 0.44(0.10,0.77) | -0.02(-0.05,0.003) | 0.37(-0.05,0.79) | -0.01(-0.04,0.02) | 0.18 |
| Age 69 | 135.0(130.2,139.7) | -0.16(-0.52,0.20) | 133.7(127.9,139.4) | -0.04(-0.40,0.33) | 0.48 |
| **DBP** |  |  |  |  |  |
| Age 36 | 75.7(73.2,78.2) | 0.03(-0.17,0.23) | 76.2(73.1,79.3) | 0.09(-0.12,0.30) | 0.69 |
| Δ 36- 53 | 0.3(0.08,0.52) | 0.003(-0.01,0.02) | 0.30(0.02,0.57) | 0.002(-0.02,0.02) | 0.93 |
| Δ 53 -69 | -0.40(-0.64,-0.17) | -0.01(-0.03,0.01) | -0.41(-0.70,-0.12) | -0.01(-0.03,0.01) | 0.42 |
| Age 69 | 74.3(71.4,77.2) | -0.07(-0.28,0.15) | 74.7(71.2,78.2) | -0.08(-0.30,0.15) | 0.68 |

**Legend:** DBP, diastolic blood pressure; log WC, natural log waist circumference; HY, hysterectomy; NM, natural menopause; SBP, systolic blood pressure. Note that BMI and WC are natural log transformed and all values presented are in log form. Δ = change per year in risk factor.

^ P value for the interaction of the association of age at period cessation with type of menopause with trajectories.

Trajectories adjusted for socioeconomic position, type of menopause, parity, time-varying hormone replacement therapy use, age at menarche, BMI at age 36 (SBP and DBP only), smoking at age 36, physical activity at age 36.

|  | **Mean trajectory for NM** | **Mean difference in trajectory per year increase in age at NM** | **Mean trajectory for HY** | **Mean difference in trajectory per year increase in HY** | **P value for interaction of type and timing of menopause** |
| --- | --- | --- | --- | --- | --- |
| **Log trig** |  |  |  |  |  |
| Age 53 | 0.31(0.26,0.36) | -0.004(-0.02,0.01) | 0.36(0.26,0.46) | -0.01(-0.03,0.001) | 0.15 |
| Δ53-69 | -0.005(-0.01,-0.001) | 0.0003(-0.0005,0.001) | -0.01(-0.01,-0.0003) | 0.0002(-0.001,0.001) | 0.63 |
| Age 69 | 0.24(0.19,0.28) | 0.001(-0.01,0.01) | 0.20(0.09,0.32) | -0.01(-0.02,0.002) | 0.32 |
| **Log HBA1c** |  |  |  |  |  |
| Age 53 | 3.64(3.63,3.66) | -0.003(-0.006,0.00004) | 3.62(3.59,3.64) | -0.002(-0.005,0.001) | 0.08 |
| Δ53-69 | 0.002(0.001,0.003) | 0.0003(0.0002,0.0005) | 0.004(0.003,0.006) | -0.00001(-0.0002,0.0002) | 0.001 |
| Age 69 | 3.67(3.66,3.69) | 0.002(-0.0005,0.005) | 3.71(3.68,3.74) | -0.002(-0.005,0.0011) | 0.042 |
| **LDL-c** |  |  |  |  |  |
| Age 53 | 3.71(3.60,3.81) | -0.02(-0.04,0.01) | 3.66(3.45,3.87) | 0.01(-0.02,0.04) | 0.34 |
| Δ53-69 | -0.02(-0.03,-0.01) | 0.0003(-0.002,0.002) | -0.03(-0.04,-0.01) | 0.0001(-0.002,0.002) | 0.95 |
| Age 69 | 3.34(3.23,3.44) | -0.01(-0.03,0.01) | 3.30(3.02,3.57) | 0.01(-0.01,0.03) | 0.24 |
| **HDL-c** |  |  |  |  |  |
| Age 53 | 1.82(1.78,1.87) | -0.002(-0.01,0.01) | 1.86(1.78,1.95) | -0.001(-0.01,0.01) | 0.93 |
| Δ53-69 | -0.003(-0.01,-0.001) | -0.0003(-0.001,0.0003) | -0.01(-0.01,-0.004) | 0.001(-0.0001,0.001) | 0.18 |
| Age 69 | 1.77(1.72,1.81) | -0.007(-0.02,0.003) | 1.67(1.58,1.77) | 0.01(-0.002,0.02) | 0.07 |

eTable 7 Unadjusted association of age at period cessation (per year increase) with blood markers from 53 to 69 years

**Legend:** HBA1c, glycated haemoglobin; HDL-c, high density lipoprotein cholesterol; HY, hysterectomy; LDL-c, low density lipoprotein cholesterol; NM, natural menopause Note that triglyceride and HBA1c are natural log transformed. Δ = change per year in risk factor.

^ P value for the interaction of the association of age at period cessation with type of menopause with trajectories.

Table 8 Association of age at period cessation (per year increase) with blood markers from 53 to 69 years, adjusted for co-variates

|  | **Mean trajectory for NM (reference for period cessation at age 50)** | **Mean difference in trajectory per year increase in age at NM** | **Mean trajectory for HY (reference for period cessation at age 50)** | **Mean difference in trajectory per year increase in HY** | **P value for interaction of type and timing of menopause** |
| --- | --- | --- | --- | --- | --- |
| **Log trig** |  |  |  |  |  |
| Age 53 | 0.46(0.31,0.60) | -0.005(-0.02,0.01) | 0.44(0.26,0.62) | -0.01(-0.02,0.005) | 0.23 |
| Δ53-69 | -0.01(-0.02,0.005) | 0.0004(-0.0004,0.001) | -0.003(-0.02,0.01) | 0.0001(-0.001,0.001) | 0.52 |
| Age 69 | 0.37(0.24,0.50) | 0.002(-0.01,0.01) | 0.41(0.23,0.59) | -0.006(-0.02,0.004) | 0.34 |
| **Log HBA1c** |  |  |  |  |  |
| Age 53 | 3.68(3.65,3.72) | -0.003(-0.005,0.0004) | 3.67(3.63,3.72) | -0.0008(-0.004,0.002) | 0.17 |
| Δ53-69 | -0.0001(-0.002,0.002) | 0.0003(0.0001,0.0005) | 0.001(-0.002,0.004) | -0.00002(-0.0002,0.0002) | 0.003 |
| Age 69 | 3.68(3.64,3.72) | 0.0024(-0.0005,0.005) | 3.69(3.65,3.74) | -0.001(-0.004,0.002) | 0.052 |
| **LDL-c** |  |  |  |  |  |
| Age 53 | 3.89(3.59,4.20) | -0.01(-0.04,0.01) | 4.03(3.65,4.41) | 0.01(-0.02,0.04) | 0.38 |
| Δ53-69 | -0.03(-0.06,-0.01) | 0.0002(-0.002,0.002) | -0.04(-0.07,-0.01) | -0.001(-0.003,0.001) | 0.85 |
| Age 69 | 3.40(3.10,3.70) | -0.01(-0.03,0.01) | 3.21(2.80,3.62) | -0.002(-0.02,0.02) | 0.36 |
| **HDL-c** |  |  |  |  |  |
| Age 53 | 1.67(1.55,1.79) | -0.002(-0.012,0.008) | 1.70(1.54,1.85) | -0.005(-0.015,0.005) | 0.75 |
| Δ53-69 | 0.006(-0.003,0.014) | -0.0003(-0.0009,0.0004) | 0.003(-0.007,0.013) | 0.0007(0.0001,0.0013) | 0.14 |
| Age 69 | 1.76(1.63,1.90) | -0.007(-0.017,0.004) | 1.72(1.55,1.88) | 0.007(-0.003,0.016) | 0.09 |

**Legend:** HBA1c, glycated haemoglobin; HDL-c, high density lipoprotein cholesterol; HY, hysterectomy; LDL-c, low density lipoprotein cholesterol; NM, natural menopause Note that triglyceride and HBA1c are natural log transformed. Δ = change per year in risk factor.

^ P value for the interaction of the association of age at period cessation with type of menopause with trajectories.

Adjusted for socioeconomic position, type of menopause, parity, time-varying hormone replacement therapy use, age at menarche, BMI at age 36, smoking at age 36, physical activity at age 36.

eTable 9 Association of type of menopause with anthropometry and blood pressure from 36 to 69 years

|  | **Unadjusted** | | | **Adjusted** | | |
| --- | --- | --- | --- | --- | --- | --- |
|  | **Mean NM (reference)** | **Association of HY** | **P*** | **Mean NM (reference)** | **Association of HY** | **P*** |
| **Log BMI** |  |  |  |  |  |  |
| Age 36 | 3.14(3.13,3.15) | 0.01(-0.02,0.04) | 0.54 | 3.16(3.12,3.19) | 0.003(-0.025,0.031) | 0.84 |
| Δ 36- 53 | 0.009(0.008,0.010) | 0.001(0.000,0.002) | 0.18 | 0.008(0.006,0.010) | 0.001(-0.0004,0.003) | 0.16 |
| Δ 53 -69 | 0.002(0.002,0.003) | -0.00003(-0.0016,0.0015) | 0.97 | 0.004(0.002,0.006) | -0.00028(-0.00186,0.00130) | 0.73 |
| Age 69 | 3.33(3.31,3.34) | 0.03(-0.01,0.06) | 0.19 | 3.35(3.30,3.40) | 0.02(-0.02,0.05) | 0.38 |
| **Log WC** |  |  |  |  |  |  |
| Age 36 | 4.31(4.30,4.32) | 0.01(-0.01,0.03) | 0.41 | 4.32(4.29,4.35) | 0.002(-0.021,0.026) | 0.84 |
| Δ 36- 53 | 0.007(0.007,0.008) | -0.0004(-0.0019,0.0012) | 0.67 | 0.007(0.005,0.009) | -0.00005 (-0.0016,0.0017) | 0.99 |
| Δ 53 -69 | 0.006(0.005,0.007) | -0.0003(-0.0019,0.0013) | 0.69 | 0.008(0.006,0.010) | -0.0005(-0.0022,0.0011) | 0.52 |
| Age 69 | 4.53(4.52,4.55) | 0.00(-0.03,0.03) | 0.95 | 4.57(4.53,4.61) | -0.01(-0.04,0.02) | 0.70 |
| **SBP** |  |  |  |  |  |  |
| Age 36 | 116.0(114.8,117.2) | -0.85(-3.51,1.81) | 0.53 | 119.0(115.5,122.4) | -0.77(-3.45, 1.90) | 0.57 |
| Δ 36- 53 | 0.90(0.79,1.01) | 0.10(-0.14,0.35) | 0.42 | 0.53(0.21,0.85) | 0.03(-0.22,0.28) | 0.80 |
| Δ 53 -69 | 0.18(0.06,0.30) | -0.09(-0.35,0.16) | 0.47 | 0.44(0.10,0.77) | -0.07(-0.33,0.19) | 0.62 |
| Age 69 | 134.1(132.5,135.8) | -0.62(-4.13,2.89) | 0.73 | 135.0(130.2,139.7) | -1.28(-4.81,2.25) | 0.48 |
| **DBP** |  |  |  |  |  |  |
| Age 36 | 74.3(73.4,75.2) | 0.61(-1.33,2.54) | 0.54 | 75.7(73.2,78.2) | 0.55(-1.39,2.50) | 0.58 |
| Δ 36- 53 | 0.41(0.33,0.48) | -0.01(-0.18,0.16) | 0.92 | 0.3(0.08,0.52) | -0.003(-0.18,0.17) | 0.97 |
| Δ 53 -69 | -0.46(-0.54,-0.38) | 0.003 (-0.17,0.18) | 0.97 | -0.40(-0.64,-0.17) | -0.002(-0.18,0.18) | 0.98 |
| Age 69 | 73.9(72.9,74.8) | 0.51(-1.60,2.63) | 0.63 | 74.3(71.4,77.2) | 0.44(-1.69,2.57) | 0.68 |

**Legend:** DBP, diastolic blood pressure; log WC, log waist circumference; HY, hysterectomy; NM, natural menopause; SBP, systolic blood pressure. Note that BMI and WC are natural log transformed and all values presented are in log form. Δ = change per year in risk factor.

*P value for association of type of menopause (HY compared with NM) with trajectory.

Adjusted for socioeconomic position, parity, age at period cessation, time-varying hormone replacement therapy use, age at menarche, BMI at age 36 (lipids, HBA1C and blood pressure only), smoking and physical activity at age 36.

**eTable 10 Association of type of menopause with blood markers from 53 to 69 years**

|  | **Unadjusted** | | | **Adjusted** | | |
| --- | --- | --- | --- | --- | --- | --- |
|  | **Mean NM (reference)** | **Association of HY** | **P*** | **Mean NM (reference)** | **Association of HY** | **P*** |
| **Log triglyceride** |  |  |  |  |  |  |
| Age 53 | 0.31(0.26,0.36) | 0.05(-0.07,0.16) | 0.41 | 0.46(0.31,0.60) | -0.02(-0.13,0.10) | 0.77 |
| Δ 53-69 | -0.005(-0.008,-0.001) | -0.002(-0.009,0.005) | 0.58 | -0.005(-0.016,0.005) | 0.003(-0.005,0.010) | 0.52 |
| Age 69 | 0.24(0.19,0.28) | 0.01(-0.08,0.11) | 0.77 | 0.37(0.24,0.50) | 0.02(-0.07,0.12) | 0.63 |
| **Log HBA1c** |  |  |  |  |  |  |
| Age 53 | 3.64(3.63,3.66) | -0.03(-0.06,0.00) | 0.06 | 3.68(3.65,3.72) | -0.01(-0.04,0.02) | 0.55 |
| Δ 53-69 | 0.002(0.001,0.003) | 0.002(0.001,0.004) | 0.01 | 0.000(-0.002,0.002) | 0.001(-0.001,0.002) | 0.37 |
| Age 69 | 3.67(3.66,3.69) | 0.01(-0.02,0.04) | 0.58 | 3.68(3.64,3.72) | 0.004 (-0.02,0.03) | 0.78 |
| **LDL-c** |  |  |  |  |  |  |
| Age 53 | 3.71(3.60,3.81) | -0.05(-0.28,0.19) | 0.69 | 3.89(3.59,4.20) | 0.13(-0.11,0.38) | 0.28 |
| Δ 53-69 | -0.02(-0.03,-0.01) | -0.002(-0.020,0.015) | 0.79 | -0.03(-0.06,-0.01) | -0.012(-0.030,0.007) | 0.21 |
| Age 69 | 3.34(3.23,3.44) | -0.09(-0.30,0.13) | 0.43 | 3.40(3.10,3.70) | -0.05(-0.26,0.16) | 0.64 |
| **HDL-c** |  |  |  |  |  |  |
| Age 53 | 1.82(1.78,1.87) | 0.04(-0.06,0.14) | 0.40 | 1.70 (1.55,1.85) | 0.03(-0.07,0.12) | 0.59 |
| Δ 53-69 | -0.003(-0.006,-0.001) | -0.0060(-0.0120,-0.0001) | 0.05 | 0.003(-0.003,0.014) | -0.003(-0.009,0.003) | 0.33 |
| Age 69 | 1.77(1.72,1.81) | -0.05(-0.15,0.04) | 0.28 | 1.72(1.65,1.88) | -0.02(-0.12,0.07) | 0.65 |

**Legend:** HBA1c, glycated haemoglobin; HDL-c, high density lipoprotein cholesterol; HY, hysterectomy; LDL-c, low density lipoprotein cholesterol; NM, natural menopause Note that triglyceride and HBA1c are natural log transformed and all values are in log form. Δ = change per year in risk factor.

*P value for association of type of menopause (HY compared with NM) with trajectory.

Adjusted for socioeconomic position, parity, age at period cessation, time-varying hormone replacement therapy use, age at menarche, BMI at age 36 (lipids, HBA1C and blood pressure only), smoking and physical activity at age 36.

1.	Cooper R, Blell M, Hardy R, et al. Validity of age at menarche self-reported in adulthood. *Journal of epidemiology and community health.* 2006;60(11):993-997.

2.	Cooper R, Mishra GD, Kuh D. Physical activity across adulthood and physical performance in midlife: findings from a British birth cohort. *American journal of preventive medicine.* 2011;41(4):376-384.
